# The CGRP receptor antagonist BIBN4096 inhibits prolonged meningeal afferent activation evoked by brief local K^+^ stimulation but not cortical spreading depression-induced afferent sensitization but not cortical spreading depression-induced afferent sensitization

**DOI:** 10.1101/151464

**Authors:** Jun Zhao, Dan Levy

## Abstract

**Introduction:** Cortical spreading depression (CSD) is believed to promote migraine headache by enhancing the activity and mechanosensitivity of trigeminal intracranial meningeal afferents. One putative mechanism underlying this afferent response involves an acute excitation of meningeal afferents by cortical efflux of K^+^ and the ensuing antidromic release of pro-inflammatory sensory neuropeptides, such as calcitonin gene-related peptide (CGRP).

**Objectives:** We sought to investigate whether (i) a brief meningeal K^+^ stimulus leads to CGRP-dependent enhancement of meningeal afferent responses, and (ii) CSD-induced meningeal afferent activation and sensitization involve CGRP receptor signaling.

**Methods:** Extracellular single-unit recording were used to record the activity of meningeal afferents in anesthetized male rats. Stimulations included a brief meningeal application of K^+^ or induction of CSD in the frontal cortex using pinprick. CSD was documented by recording changes in cerebral blood flow using laser Doppler flowmetery. CGRP receptor activity was inhibited with BIBN4096 (333μM, i.v.).

**Results:** Meningeal K^+^ stimulation acutely activated 86% of the afferent tested and also promoted in ~65% of the afferents a 3-fold increase in ongoing activity which was delayed by 23.3±4.1 min and lasted for 22.2±5.6 min. K^+^ stimulation did not promote mechanical sensitization. Pretreatment with BIBN4096 suppressed the K^+^- induced delayed afferent activation, reduced CSD-evoked cortical hyperemia, but had no effect on the enhanced activation or mechanical sensitization of meningeal afferents following CSD.

**Conclusion:** While CGRP-mediated activation of meningeal afferents evoked by cortical efflux of K^+^ could promote headache, acute activation of CGRP receptors may not play a key role in mediating CSD-evoked headache.

**Previous presentation of the research, manuscript, or abstract:** Parts of the manuscript have been presented previously only in an abstract form.

## 1. Introduction

Migraine is the third most prevalent and seventh most disabling disease in the world, affecting about 15% of the adult population worldwide [60,62]. A key migraine theory links the genesis of the headache to a cascade of inflammatory-driven nociceptive events, namely the prolonged activation and increased mechanosensitivity of pain sensitive afferents that innervate the intracranial meninges and their related large blood vessels [36,42,47,48].

The origin of such meningeal inflammatory response in migraine is not completely understood, although a key hypothesis implicates the action of proinflammatory sensory neuropeptides that are released from peripheral nerve endings of activated primary afferents through an “axon reflex” process [29,46]. According to this neurogenic inflammatory hypothesis, the initial stimulation of trigeminal meningeal afferents leads to antidromic calcium-mediated vesicular release of sensory neuropeptides, which act upon the meningeal vasculature and immune cells to promote sterile inflammation that subsequently leads to the prolonged increase in meningeal afferent responsiveness [7,9,36,52]. While this hypothesis was initially focused on the neuropeptide substance P as the key mediator of meningeal inflammation and migraine pain [52], the failure of substance P receptor antagonists to abort migraine attacks [10] has led to diminished support for the role of neurogenic inflammation in migraine headache origin.

Another key neuropeptide that is released from activated trigeminal afferents is calcitonin gene-related peptide (CGRP). CGRP is a potent vasodilator of cerebral [41] and dural arteries [33,43] and may also contribute to dural plasma extravasation [28]. Trigeminal CGRP is considered a major player in migraine pain: a notion that was based primarily on a study which detected elevated CGRP levels in the jugular vein during a migraine attack and its normalization by the anti-migraine drug sumatriptan [25]. A link between CGRP, meningeal axonal reflex and migraine headache has been further argued based on studies showing that agents that block CGRP action and inhibit neurogenic meningeal vasodilatation [51,65,69] can abort migraine headache [1,31]. The notion that meningeal elaboration of CGRP can directly enhance the responsiveness of meningeal afferents, however, has been questioned by us previously based on data from experiments that tested the effects of exogenously applied CGRP [37]. Nonetheless, the notion that anti-migraine treatments that block CGRP action, including drugs from the gepant class of CGRP receptor antagonists and monoclonal antibodies, act peripherally [15] suggests that neurogenic release of CGRP does, somehow, play a role in mediating migraine pain.

One key unanswered question is what process leads to the initial trigeminal peripheral release of CGRP in migraine. One such potential event might be cortical spreading depression (CSD), a putative trigger of migraine with aura [34]. CSD is associated with a robust increase in cerebral blood flow [49], which is mediated in part by cerebrovascular release of CGRP [8,55,67]. A more direct link between CSD and the origin of migraine headache has been suggested by our previous studies in a rat model, which documented in a substantial number of meningeal afferents a short-lasting increase in ongoing activity, during the passage of the CSD wave under the afferents’ receptive field that is followed, after some delay, by a prolonged activation and mechanosensitization of the afferents [71,74,75]. The mechanism underlying the short-lasting activation of meningeal afferents during the CSD event remains unknown but may involve the nociceptive action of diffusible excitatory molecules, including potassium ions [63] that are briefly released in the cortex during the CSD wave [4,20,40,66]. Based on the neurogenic inflammatory hypothesis of migraine in the wake of CSD, the initial activation of meningeal afferents by K^+^ or potentially other nociceptive factors, indirectly leads to the prolonged and enhanced afferent responses through a mechanism that involves a trigeminal axon reflex and meningeal release of CGRP or other proinflammatory neuropeptides [52].

Here, we used single-unit recording of meningeal afferents in anesthetized rats to initially address the question, whether a brief excitation of meningeal afferents in response to short-term elaboration of K^+^, as would be expected during CSD, leads to prolonged change in the afferents’ response properties. Having identified such K^+^-driven afferent response, and given that acute exposure of meningeal afferents to K^+^ likely promotes meningeal CGRP release [11,12,43], we further examined whether acute CGRP-receptor signaling mediates this response. Finally, given that CSD is associated with a CGRP-related meningeal vascular response we investigated whether CGRP receptor signaling contributes to the prolonged meningeal afferent response following CSD.

## 2. Materials and Methods

### 2.1 Animals and anesthesia

The current study employed male Sprague-Dawley rats (250–350 g). Animals were handled in compliance with the experimental protocol approved by the Institutional Animal Care and Use Committee of the Beth Israel Deaconess Medical Center. Animals were deeply anesthetized with urethane (1.5 g/kg, ip) and mounted on a stereotaxic frame (Kopf Instruments). Core temperature was kept at 37.5-38°C using a homoeothermic control system. Animals breathed spontaneously room air enriched with 100% O2. Physiological parameters were collected throughout the experiments using PhysioSuite^®^ (Kent Scientific) and CapStar-100 (CWE). Data used in this report were obtained only from animals exhibiting physiological levels of oxygen saturation (>95%), heart rate (350–450 bpm), and end-tidal CO2 (3.5–4.5%).

### 2.2. Surgery and electrophysiological recordings

A saline-cooled dental drill was used to perform a craniotomy to expose the left transverse sinus as well as the adjacent cranial dura extending ∼2 mm rostral to the sinus. In animals tested for the effects of CSD, a small burr hole (0.5-mm diameter) was drilled to expose a small area of dura above the frontal cortex to induce CSD [75]. The exposed dura was bathed with a modified synthetic interstitial fluid (SIF) containing 135 mM NaCl, 5 mM KCl, 1 mM MgCl2, 5 mM CaCl2, 10 mM glucose, and 10 mM HEPES, pH 7.2. The femoral vein was cannulated to allow for intravenous administration. Single-unit activity of meningeal afferents (1 afferent/rat) was recorded in the ipsilateral (left) trigeminal ganglion using a contralateral approach, as described recently [73,74] (and see also Figure 1). Briefly, a small craniotomy (2 × 2 mm) was made in the calvaria over the right hemisphere, 2 mm caudal and 2 mm lateral to Bregma. A platinum-coated tungsten microelectrode (impedance 100 kΩ, FHC) was advanced through the right hemisphere into the left trigeminal ganglion using an angled (22.5°) trajectory. In earlier studies, we verified that inserting the recording electrode using this approach did not produce CSD in the left cortical hemisphere. Meningeal afferents were identified by their constant latency response to single shock stimulation applied to the dura above the ipsilateral transverse sinus (0.5-ms pulse, 5 mA, 0.5 Hz). The response latency was used to calculate conduction velocity (CV), based on a conduction distance to the trigeminal ganglion of 12.5 mm [63]. Neurons were classified as either A-delta (1.5 < CV ≤ 5 m/s) or C-afferents (CV ≤ 1.5 m/sec). Neural activity was digitized and a real-time waveform discriminator (Spike 2 software, CED) was used to create and store a template for the action potential evoked by electrical stimulation, which was used later to acquire and analyze the ongoing activity of the neurons and the activity evoked by mechanical stimulation and CSD.

**Figure 1:**
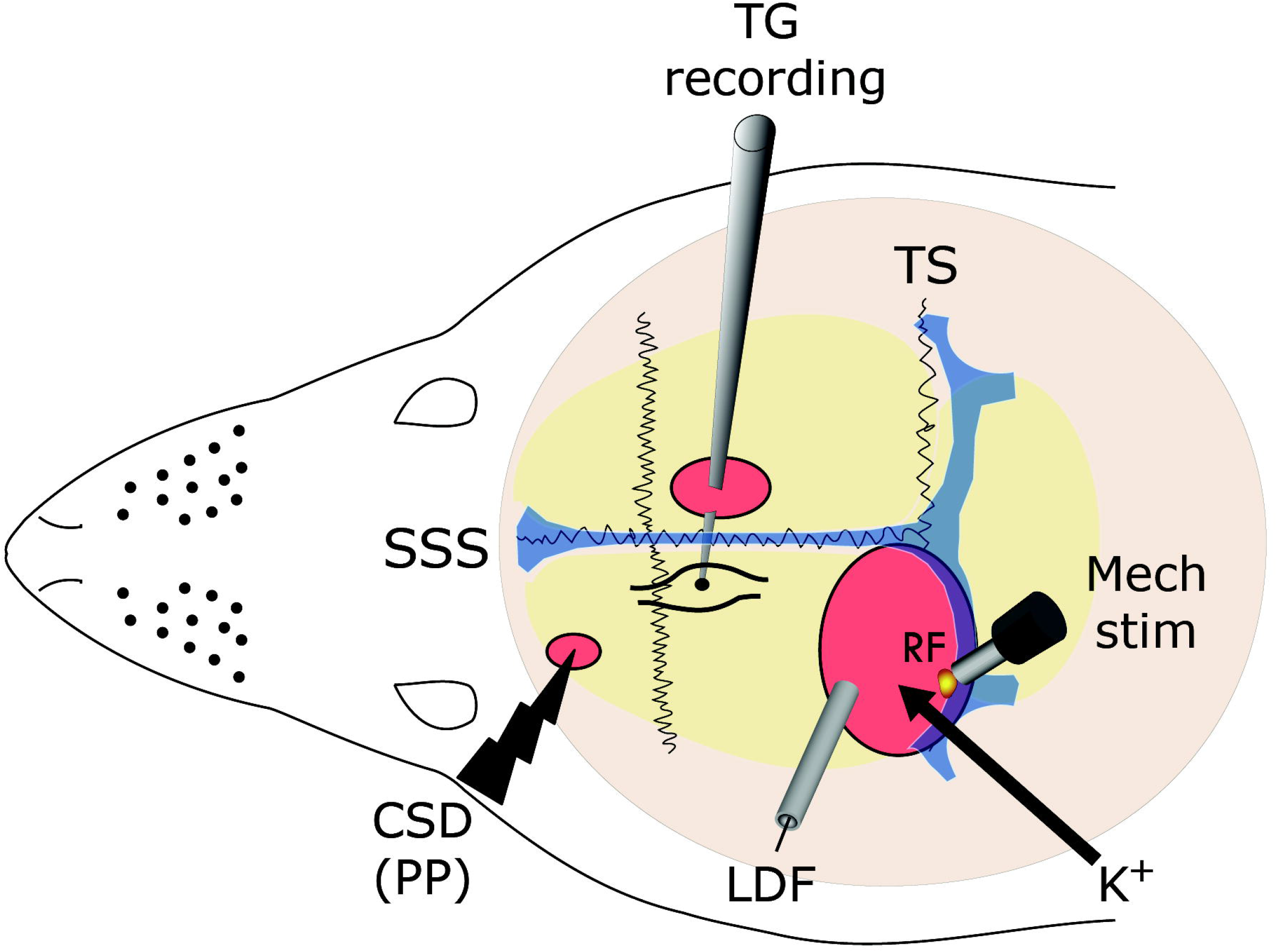
Experimental setup: Three skull openings (red ovals) were made. A small burr hole was made over the left frontal cortex to elicit a single cortical spreading depression (CSD) event using a pinprick (PP) stimulation, by inserting a sharp glass microelectrode 2mm into the cortex. Meningeal afferent activity was recorded in the left trigeminal ganglion (TG) using a tungsten microelectrode inserted through a craniotomy made over the contralateral hemisphere. An ipsilateral craniotomy was made to expose a part of the left transverse sinus (TS) and its vicinity to search for meningeal afferents with mechanical receptive field (RF). Quantitative mechanical stimuli were delivered to the afferents RF using a feedback-controlled mechanical stimulator. Laser Doppler flowmetry (LDF) probe was placed over the cortex near the stimulated afferents RF to validate the induction of the CSD by testing related changes in cerebral blood flow. SSS, superior sagittal sinus. K^+^ stimulation was made by applying KCl to the exposed dural RF of the recorded afferent.

### 2.2 Mechanical stimulation and detection of sensitization

Mechanical receptive fields (RF) of meningeal afferents were first identified by probing the exposed dura with a 6.9 g von Frey filament (Stoelting). The site of lowest mechanical threshold was further determined using monofilaments that exert lower forces (>0.03 g). For quantitative determination of mechanical responsiveness, we applied mechanical stimuli to the site with the lowest threshold, using a servo force-controlled mechanical stimulator (Series 300B Dual Mode Servo System, Aurora Scientific). The stimulus was delivered using a flat-ended cylindrical plastic probe that was attached to the tip of the stimulator arm. One of three probe diameters (0.5, 0.8 or 1.1 mm) was selected for each neuron, depending on the sensitivity of the neuron. Stimulus trials for testing mechanically sensitive were made using a custom-written script for Spike 2 and consisted of ramp and hold stimuli (rise time 100 ms, stimulus width 2 s, inter-stimulus interval 120 s). Each trial included a threshold stimulus (TH, which normally evoked 1-3-Hz responses) followed by a supra-threshold stimulus (STH, usually X2 of the threshold pressure; 8-10 Hz responses). To minimize response desensitization, stimulus trials were delivered every 15 min throughout the experiment [70,75]. Ongoing afferent discharge rate was recorded continuously between the stimulation trials. Responses to mechanical stimulation were determined during at least four consecutive trials before the elicitation of CSD. Only afferents that exhibited consistent responses (variation of <0.5 Hz for TH responses and <1.5 Hz for STH responses) during baseline recordings were tested further [75].

### 2.3 Induction and monitoring of CSD

A single CSD episode was induced in the frontal cortex, using a pinprick stimulus with a fine glass micropipette (diameter 10 μm) at ∼2 mm depth for 2 sec. CSD was induced in the frontal cortex to avoid potential damage to the meningeal tissue near the tested RF of the studied afferents, which could have led to their activation and/or sensitization [74]. The occurrence of a CSD episode was determined noninvasively by recording changes in cerebral blood flow (CBF) using a laser Doppler flowmeter (LDF, Laserflo, Vasamedics) with the needle probe (0.8 mm diameter) positioned ~1 mm above the exposed dura, 1-2 mm rostral to the RF of the recorded afferents (Figure 1) in an area devoid of large blood vessels (>100 μm). Such LDF system (using a near IR 785 nm probe and fiber separation of 0.25 mm) is expected to record CBF changes at a cortical depth of ~1 mm [21]. LDF recordings were obtained with ambient lights turned off. CBF data was digitized and continuously sampled via an A/D interface (power 1401 and spike 2 software, CED, Cambridge, UK). Induction of CSD was considered successful when the typical hemodynamic signature characterized by a large transient (∼1-2 min) cerebral hyperemia, followed by prolonged (>1h) post-CSD oligemia was observed [24,53,74].

### 2.4 Drugs

BIBN4096BS (Tocris; Cat. No. 4561) was dissolved in DMSO to make a 50mM stock solution. For intravenous injection, a 400μM solution was made by diluting with 0.9% saline. Potassium chloride (Sigma Aldrich; Cat. No. P9333) stock solution (1M) was made in distilled water and further diluted to 50mM using SIF. For local meningeal K^+^ stimulation, we used a piece of cotton wool to create a circular pool to cover the exposed dural RF of the recorded afferent. The K^+^ stimulation protocol consisted of an initial application of 20μl of a 50mM KCl solution for 10 seconds, followed by 5 sequential applications of 20μl of SIF (which contains 5mM K^+^) every 10 sec and a final wash using 500μl of SIF.

### 2.5 Data analysis

Offline analyses for afferent responses were conducted using template matching in Spike 2. Statistical analyses were conducted using Prism 7 software. Average data are presented in the text as the mean ± SEM. Average data in figures is presented as the mean ± 95% confidence interval (CI). Criteria used to consider meningeal afferent activation and sensitization responses were based on our previous studies [74,75]. In brief, an increase in afferents ongoing activity was considered if the firing rate increased above the upper end point of the 95% CI calculated for the baseline mean. Acute afferent activation was considered when ongoing activity level increased during the K^+^ stimulus or the arrival of the CSD wave (i.e. the hyperemia stage) under the RF of the studied afferent and subsided within 2 minutes. Prolonged activation was considered when ongoing activity rate was increased for >10 min. Mechanical sensitization was considered only if the afferent responses changed according to the following criteria: TH and/or STH responses increased to a level greater than the upper endpoint of the 95% CI calculated for the baseline mean; increased responsiveness began during the first 60 min post-CSD; and sensitization lasted for at least 30 min. For analyses of CSD-related changes in the afferent responses, data from A-delta and C afferents were combined given their comparable values. Afferent responses to CSD in the presence of BIBN4096BS were compared to a pool of data from our earlier vehicle studies [74,75] that was combined with additional data obtained from new vehicle experiments (total n=36) conducted to increase sample size and add power. Group differences were analyzed using two-tailed, Fisher’s exact test. Statistical differences were analyzed using two-tailed unpaired *t*-test or Mann Whitney U test for datasets that failed normality test. Results were considered to be significant at *p*<0.05.

## 3. Results

### 3.1 Brief meningeal stimulation with a CSD-related K^+^ stimulus promotes prolonged activation of meningeal afferents

Earlier studies suggested that in response to CSD the level of extracellular parenchymal K^+^ rises quickly within the superficial cortex to ~50mM followed by a rapid decrease within ~ 1 minute to baseline levels [20,32,40,45,66]. We therefore first asked whether a brief, local meningeal stimulation with a similar pattern of changes in K^+^ concentration (See Figure 2B) could promote long-term changes in the activity and/or mechanosensitivity of meningeal afferents, as observed in some afferents following CSD [71,74]. The effect of K^+^ stimulation was tested on 14 meningeal afferents (5 A-delta, 9 C). Local administration of K^+^ gave rise, as expected [63], to a robust yet short-term (< 70 sec) increase in the ongoing activity rate (8.7±2.4 fold) in 12 afferents (4/5 A-delta, 8/9 C). Following a wash with SIF (which contains 5mM K^+^), 9/14 of the afferents (3/5 A-delta 6/9 C) also developed a delayed and prolonged increase in their ongoing activity rate (see an example of the immediate and delayed activation of a C unit in Figure 2A). The responses of the A delta and C afferents were similar and therefore combined for further analyses. As Figure 3 depicts, the prolonged afferent activation following the K^+^ stimulus developed with an average delay of 23.3±4.1 min (range 5-40 min) and lasted for 22.2±5.6 min (range 10-55 min). Overall, the K^+^-stimulation evoked a 3.1±0.4 fold (range 1.7-5.0 fold) increase in the afferent firing rate. When compared to the CSD-evoked prolonged meningeal afferent activation recorded in animals treated with vehicle (9/19 A-delta, 9/17 C), the afferent activation induced by the K^+^ stimulation developed with a longer delay (compare to onset of 8.8±2.6 min; range 0-35 min in the CSD group; p<0.05, unpaired *t*-test). The duration of the prolonged afferent activation following the K^+^ stimulation was also shorter (compare to duration of 32.8±3.8 min; range 10-55 min in the CSD group; p<0.05, *t*-test). The magnitude of the prolonged response following K^+^ stimulation, however, was not different between the two groups (compare to 2.9±0.5 fold; range 1.2-10.4 fold in the CSD group; p=0.21, Mann Whitney test). As Figure 4 illustrates, following K^+^ stimulation meningeal afferents did not become sensitized to mechanical stimulation. We detected a brief (30 min) increased in responsiveness in only 2/24 C afferent, which was not different that that observed in time-control experiments (0/6 A-delta and 1/6 C showed increased responsiveness for 30 min).

**Figure 2.**
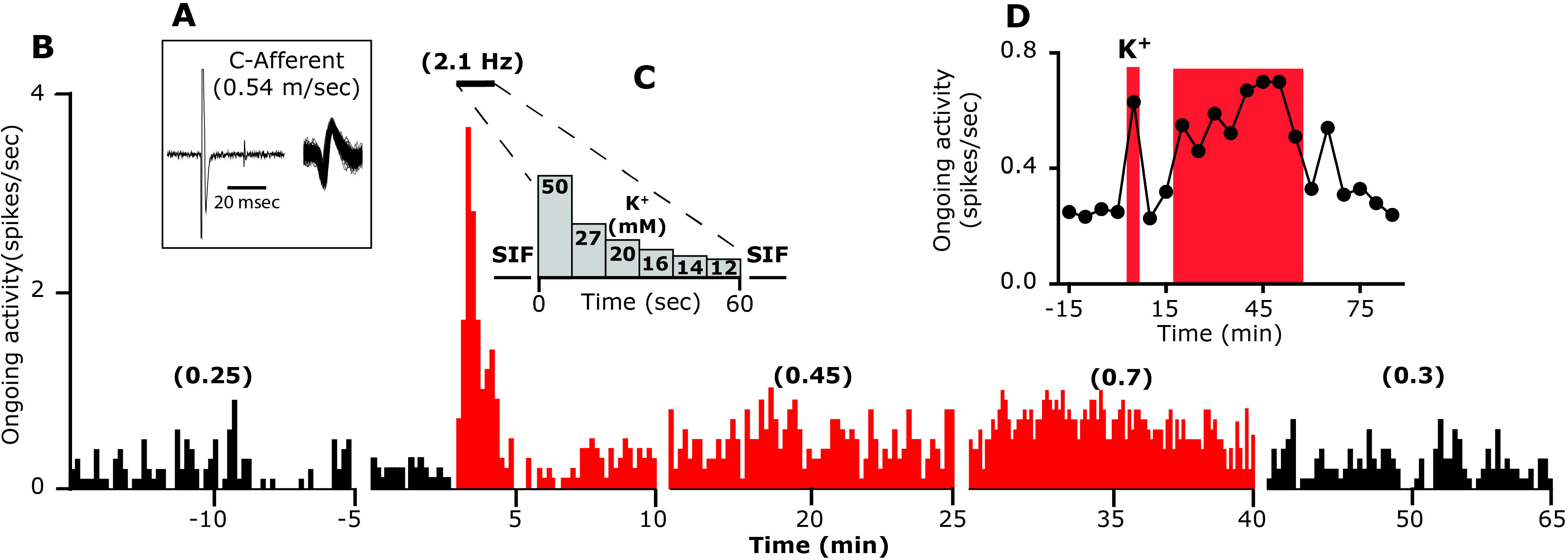
An example showing the development of delayed and prolonged activation following acute local stimulation with K^+^ in one C meningeal afferent. **A.** The afferent response latency to electrical stimulation used to calculate the conduction velocity is shown on the left. An overdrawn spike waveform of the afferent used for data analysis is shown on the right. **B.** Traces represent peristimulus time histograms (PSTH, 10-min epochs, 10-sec bin-size). Average ongoing activity rate (spikes/sec) is denoted in parentheses. The last 5 min recording period during the mechano-stimulation trials, conducted at the end of each 15-min recording epoch, were omitted for clarity. **C.** The protocol employed for meningeal application of K^+^. Note the acute increase in afferent activity during the stimulation. **D.** Time course data depicting the ongoing activity level of the same afferent depicted in panel B during baseline and every 5 min after K^+^ stimulation. Red color areas represent the acute and delayed phases of neuronal activation.

**Figure 3.**
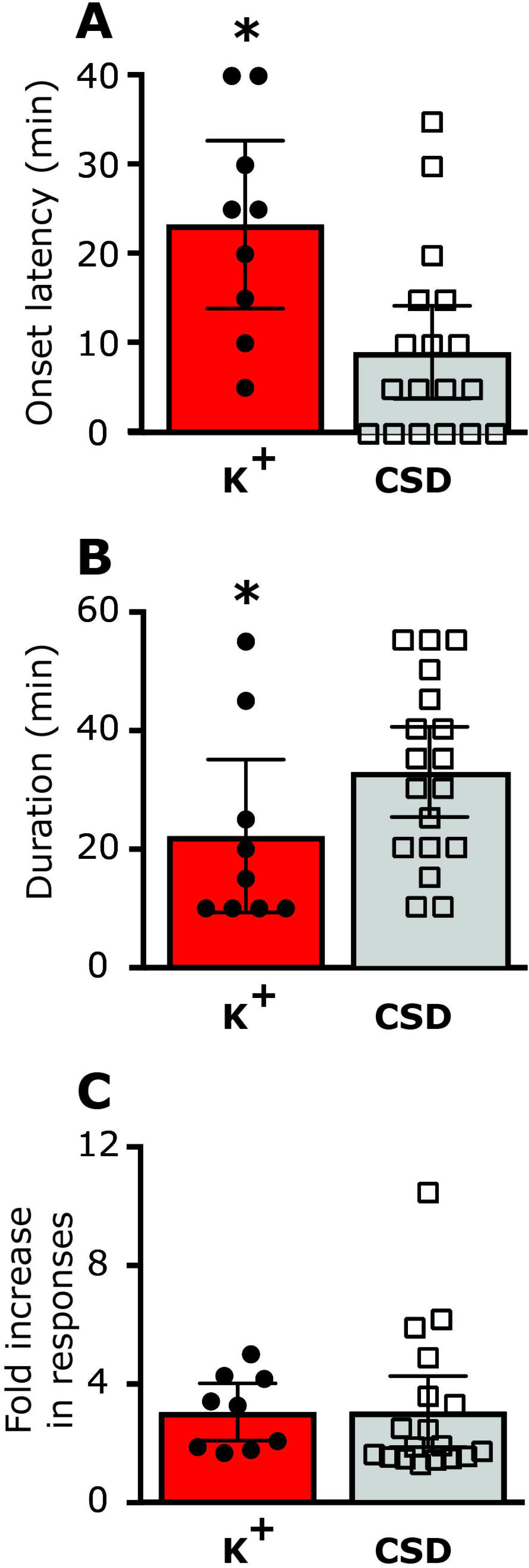
Characteristics of the delayed and prolonged meningeal afferent responses induced by local K^+^ stimulation compared to that induced by CSD in vehicle treated animals. **A**, onset latency. **B**, Duration of the afferent activation. **C**, Magnitude of the increase in ongoing activity. * p<0.05; unpaired *t*-test.

**Figure 4.**
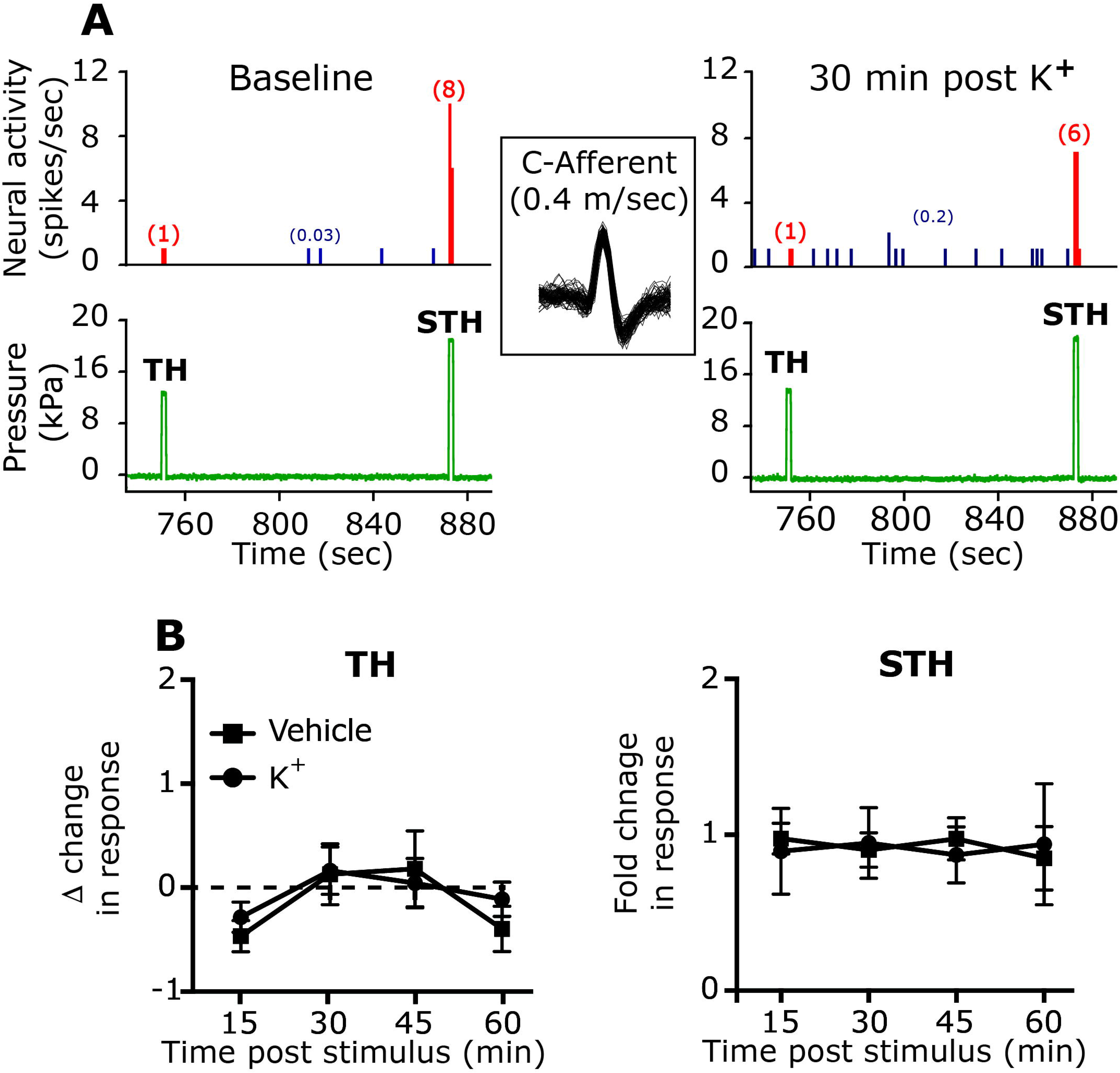
Meningeal K^+^ stimulus does not affect the mechanical responsiveness of meningeal afferents. **A.** An example of two experimental trials depicting the responses of one C afferent (an overdrawn spike waveform is depicted in the insert) to threshold (TH) and suprathreshold (STH) mechanical stimuli (green traces) applied to the its dural receptive field during baseline recording and then at 30 min following local K^+^ stimulation. Red numbers in parentheses denote mechanically-evoked afferent responses (spikes/sec). Note the lack of changes in TH or STH responses. Blue numbers in parentheses denote ongoing activity rate, note the K+ evoked activation. **B.** Summary of time course data collected from meningeal afferents in time control studies (vehicle) and following local K^+^ stimulation showing lack of changes in TH and STH responses.

### 3.2 K^+^ - induced prolonged activation of meningeal afferents is CGRP-dependent

Previous studies reported that a brief meningeal stimulation with high concentration of K^+^ similar to the current study evokes meningeal CGRP release [12,43,58]. We therefore asked whether the K^+^ - driven prolonged activation of meningeal afferents is CGRP-dependent, and thus potentially related to meningeal afferent axonal reflex. To examine the role of CGRP signaling we administered the CGRP-receptor antagonist BIBN4096 (333 μg/kg, i.v) one hour before testing the afferent response to meningeal K^+^ stimulation. The dose of BIBN4096 was adapted from previous studies showing efficacy in inhibiting neurogenically-evoked meningeal vasodilation [26,51,65] and meningeal afferent volleys in the spinal trigeminal nucleus [22]. We first tested the effect of BIBN4096 on the baseline response properties of meningeal afferents in 36 afferents (19 A-delta, 18 C). When compared to data obtained from animals administered only with saline (50 afferents; 25 A-delta, 25 C), BIBN4096 treatment neither affected the afferents baseline ongoing activity rate (Table 1), nor influenced their baseline mechanosensitivity (Table 2). We next tested the effect of BIBN4096 treatment on the K^+^ -evoked afferent responses in 6 A-delta and 7 C afferents. BIBN4096 did not inhibit the initial short excitatory response to K^+^ (10/13; 4/6 A-delta; 6/7 C afferents developed a 7.6±3.7 fold increase in activity, p=0.64, t-test). However, as Figure 5 depicts, in the presence of BIBN4096 only 2/13 (2 C) of the afferents developed a prolonged increase in ongoing activity (compared with 9/14 units affected in vehicle control experiments, p<0.05; two-tailed, Fisher’s exact test). The onset latencies for the prolonged activation in these two afferents were 35 and 10 min with matching durations of 20 and 10 min. The magnitude of change in afferent activity in animals administered with BIBN4096 prior to K^+^ stimulation was also lower than that observed in units treated with vehicle prior to K^+^ stimulation (p<0.05, t-test).

**Table 1:** Systemic BIBN4096 treatment does not affect baseline ongoing activity of meningeal afferents.

|  | Before treatment |  | 60 min post-treatment |  |
| --- | --- | --- | --- | --- |
|  | A-delta | C | A-delta | C |
| **Vehicle** | 0.7±0.1 | 1.1±0.2 | 0.6±0.1 | 1.0±0.2 |
| **BIBN4096** | 0.5±0.1 | 1.2±0.3 | 0.5±0.3 | 1.1±0.3 |

Data is expressed as the mean rate of ongoing activity (spikes/sec) ± SEM of A-delta and C meningeal afferents recorded from the TG of animals before systemic administration of BIBN4096 (19 A-delta, 18 C) or saline (25 A-delta, 25 C) and 60 min later (post-treatment), prior to meningeal K<sup>+</sup> stimulation or induction of CSD. Data analyzed using paired t-test showed no effect of treatments. TG, trigeminal ganglion.

Data is expressed as the mean rate of ongoing activity (spikes/sec) ± SEM of A-delta and C meningeal afferents recorded from the TG of animals before systemic administration of BIBN4096 (19 A-delta, 18 C) or its vehicle (25 A-delta, 25 C) and 60 min later (post-treatment), prior to meningeal K^+^ stimulation or induction of CSD. Data were analyzed using paired t-test and showed no effect of treatments. TG, trigeminal ganglion.

**Table 2:** Systemic BIBN4096 treatment does not affect baseline mechanosensitivity of meningeal afferents.

|  | Pre-treatment |  | 60 min post-treatment |  |
| --- | --- | --- | --- | --- |
|  | TH | STH | TH | STH |
| **Control** | 1.3±0.3 | 8.2±1.0 | 2.0±0.7 | 8.3±0.9 |
| **BIBN4096** | 2.1±0.3 | 8.7±0.9 | 2.8±0.6 | 9.9±1.2 |

Data is expressed as the mean rate of neural responses (spikes/sec) ± SEM of meningeal afferents evoked during the 2 sec mechanical stimuli of Threshold (TH) and suprathreshold (STH) forces applied to afferents dural receptive field prior to, and following systemic treatment with BIBN4096 (n=33) or its vehicle (n=28). Data were analyzed using paired t-test and showed no effect of treatments.

**Figure 5.**
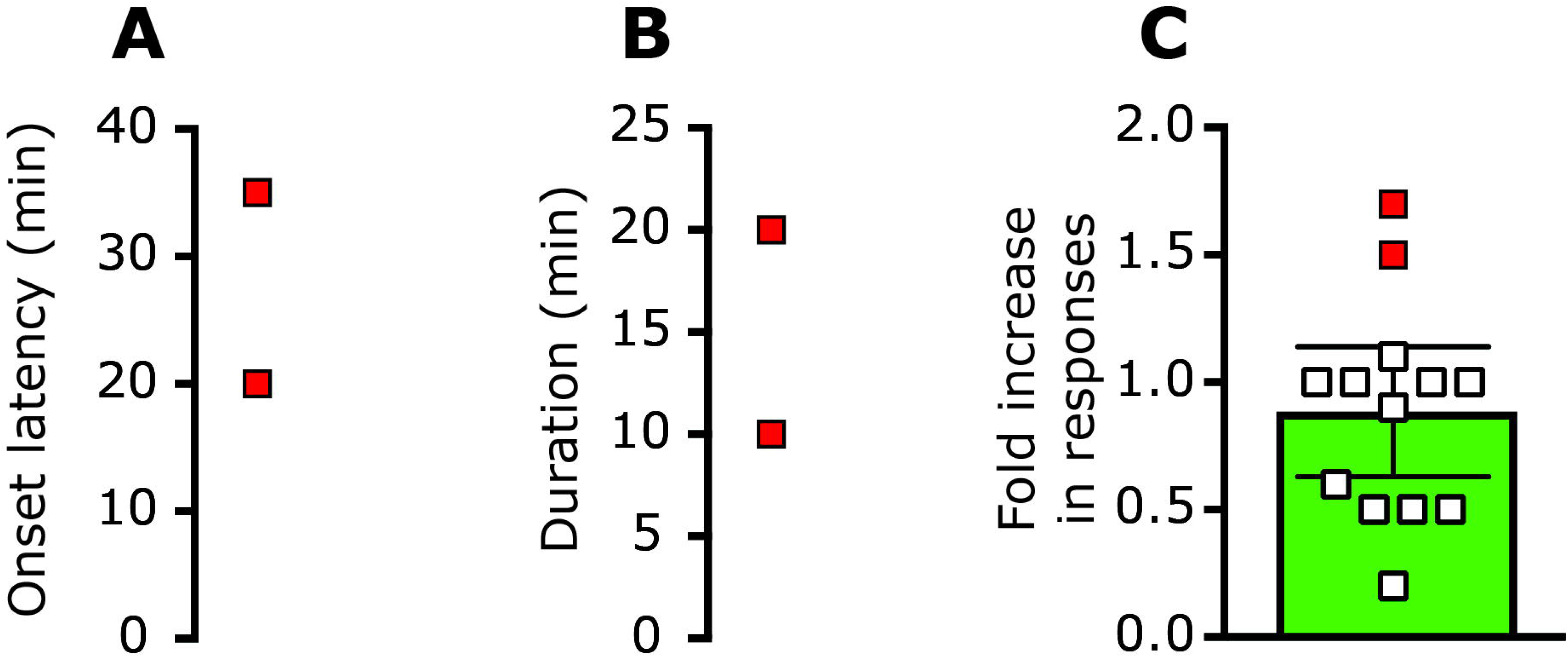
Inhibition of K^+^ -induced prolonged meningeal afferent activation by BIBN4096. In the presence of BIBN4096, only 2 afferents became activated (red symbols). **A.** onset latency. **B.** Duration of the afferent activation. **C.** Magnitude of change in ongoing activity rate. Open symbols denote changes in the activity of afferents not activated following K^+^ stimulation in BIBN4096 pretreated animals.

### 3.3 Enhanced meningeal afferent activation following CSD is not mediated by acute CGRP receptor signaling

Given the ability of BIB4096 pretreatment to inhibit the K^+^ - driven prolonged afferent activation, we next tested whether CGRP receptor activation also contributes to the prolonged meningeal afferent’s responses in the wake of CSD. We initially verified that in the setting of CSD BIBN4096 treatment inhibits a previously recognized CGRP-mediated meningeal response, namely the brief pial hyperperfusion [8,67]. Systemic administration of BIBN4096 (n=15) did not affect baseline CBF (Figure 6A) but inhibited the cortical hyperemic response during CSD by ~25% (Figure 6B). Induction of CSD in BIBN4096 pretreated animals gave rise to a hyperemic response that was significantly lower than that observed in vehicle-treated animals (1.56±0.1 fold vs 2.1±0.1 fold; p<0.01, Mann Whitney test) confirming the efficacy of BIBN4096 as a CGRP receptor antagonist in this model. The effect of BIBN409 administration on CSD-evoked meningeal afferent activation was tested in 24 afferents (13 A-delta and 11 C). In the presence of BIBN4096, CSD resulted in the development of prolonged activation in 9/24 (37.5%) of the afferents tested (4/13 A-delta; 5/11 C). This response rate was not different than that observed in afferents recorded in animals treated only with vehicle (18/36; p=0.80 overall; p=0.47 for A-delta afferent population; p=1.00 for the C-afferent population, Fisher’s exact test). BIBN4096 treatment also did not influence the characteristics of the CSD-evoked prolonged afferent activation. The average onset latency for CSD-evoked afferent activation in the BIBN treated group (3.9±1.6 min; range 0-15 min; Figure 7A) was not different than that observed in the vehicle treated group (p=0.29, t-test, Figure 3A). The duration of the post-CSD afferent discharge in the BIBN4096 treatment group (average 30.0±5.3 min; range 10-55 min; Figure 7B) was also not different than that observed in the vehicle control group (p=0.15; t-test, Figure 3B). Finally, in the presence BIB4096, the magnitude of the increase in afferent discharge rate following CSD (average 2.9±0.7 fold; range 1.5-8.3 fold; Figure 7C) was also not different when compared to that observed in the vehicle treatment group (p=0.41; t-test, Figure 3C).

**Figure 6.**
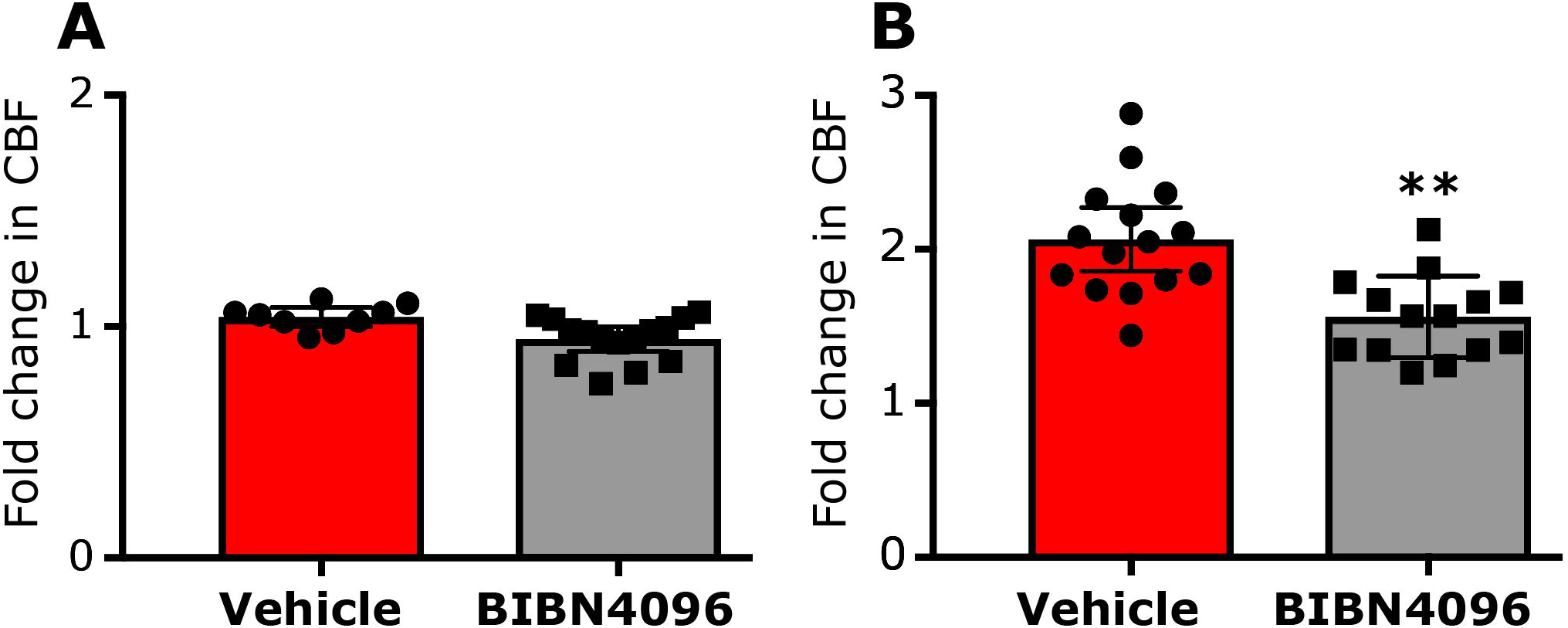
The effect of BIBN4096 on CSD-evoked cerebrovascular hyperemia. **A.** Changes in baseline cerebrovascular perfusion. **B.** Cerebrovascular dilation during CSD.

**Figure 7.**
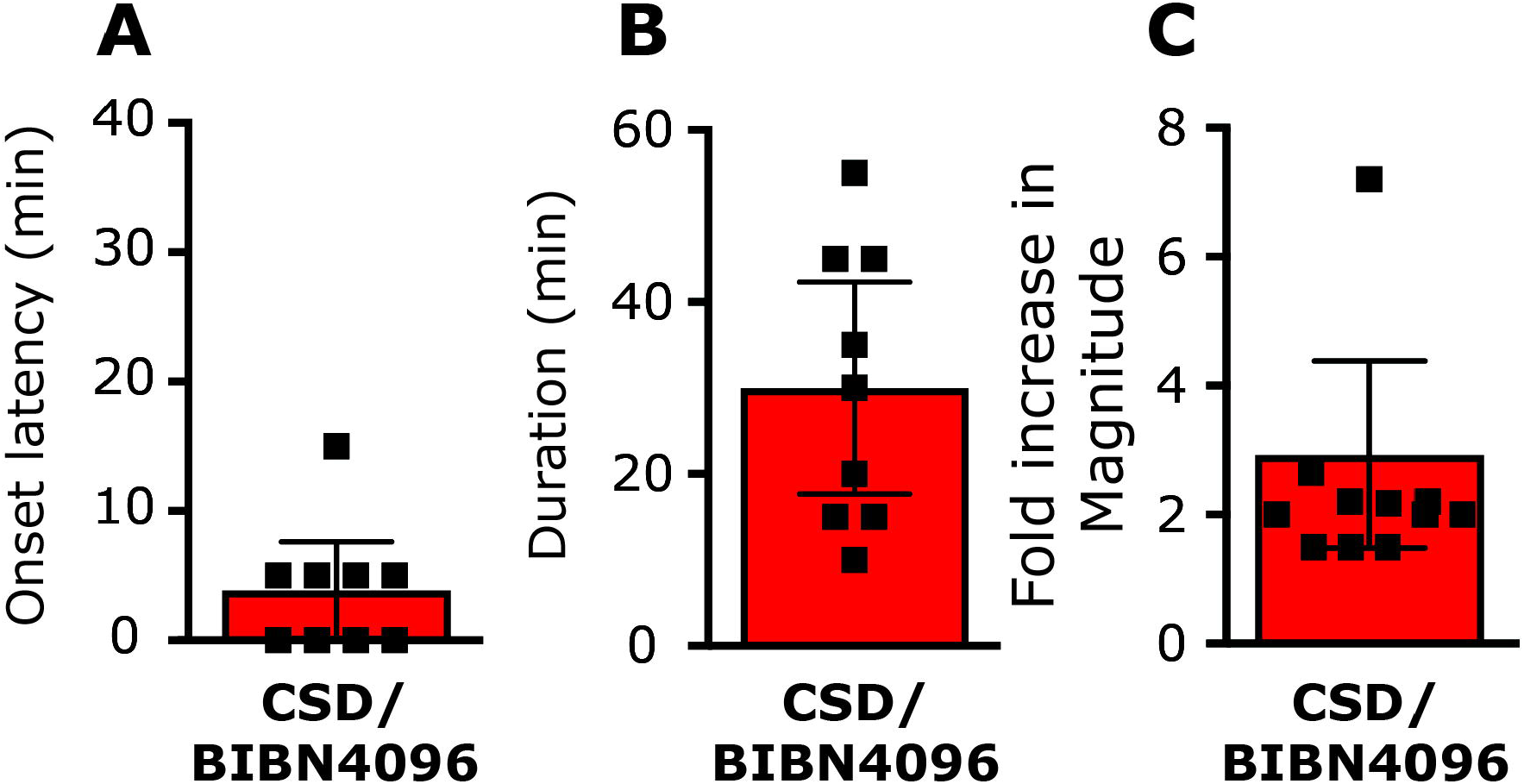
The effect of BIBN4096 on CSD-evoked prolonged activation of meningeal afferents. Onset latency **(A)**, duration **(B)**, and magnitude **(C)** of the prolonged increase in ongoing activity of meningeal afferents following CSD in animals pretreated with BIBN4096.

### 3.4 Acute CGRP receptor signaling does not mediate CSD-evoked mechanical sensitization of meningeal afferents

To examine the effect of BIB4096 pretreatment on the CSD-evoked meningeal afferent mechanosensitization we analyzed data collected from 21 afferents (11 A-delta and 10 C), all of which displayed consistent responses at baseline. In BIBN4096-treated animals, CSD evoked mechanical sensitization in 11/21 afferents (5/11 A-delta and 6/10 C, see Figures 8A,B). This rate of response was not different than that observed in animals treated with vehicle (7/10 A-delta and 9/13 C; p=0.35 overall; p=0.39 for the A-delta afferent population; p=0.69 for the C afferent population, Fisher’s exact test). The magnitudes of the TH and STH sensitization responses in the BIBN treated group were not different than those observed in the control group (TH, Δ1.6±0.5 spikes/sec; range 0.3-4.8 spikes/sec vs. Δ1.8±0.4 spikes/sec; range 0.3-4.8 spikes/sec, p=0.74 Mann Whitney test; STH, 1.6±0.1 fold; range 1.2-2.3 fold vs. 1.6±0.1 fold; range 1.2-2.0 fold; p=0.92; Mann Whitney test). BIBN4096 also did not affect the onset latency for the CSD-evoked sensitization. In the BIB4096-treated group TH sensitization latency averaged 35.6±6.3 min (range 15-60 min) and was not statistically different than that observed in the control group (21.0±3.6 min, range 15-45 min; p=0.09; Mann Whitney test; Figure 8C). The onset latency for the STH sensitization in the BIBN4096 group was 31.5±5.7 min (range 15-60), which was also not statistically different than that observed in the control group (24.2±3.2 min; range 15-60 min; p=0.25; Mann Whitney test; Figure 8D). The duration of the TH sensitization response in the BIBN4096-treated group was 73.8±9.9 min (range 30-120 min) and was also similar to that observed in the control group (71.3±10.5 min; range 30-105 min, p=0.89; Mann Whitney test; Figure 8E). Finally, the duration of the STH sensitization response in the BIBN4096 group was 64.5±9.8 min (range 30-120 min) and was not statistically different than that observed in the control group (73.9±8.7 min; range 30-120 min; p=0.54; Mann Whitney test; Figure 8F).

**Figure 8.**
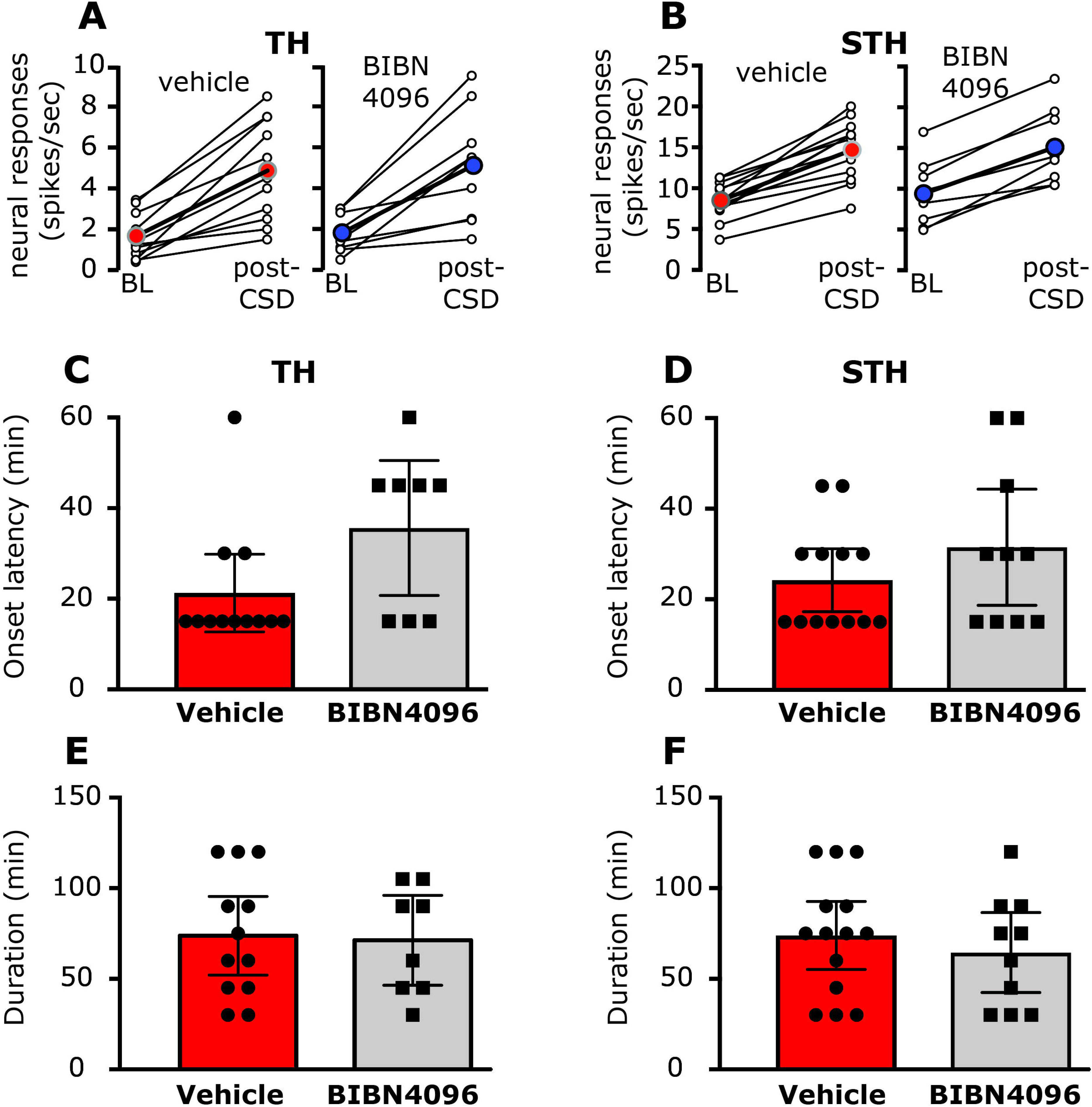
BIBN4096 does not inhibit CSD-induced mechanical sensitization of meningeal afferents. TH **(A)** and STH **(B)** responses in neurons that exhibited mechanical sensitization in response to CSD following treatment with vehicle or BIBN4096. Open circles depict the responses at baseline, before CSD, and during the time of peak response after CSD of individual afferents. Colored circles denote the mean response for the population. Note the similar magnitude of changes in the vehicle and BIBN4096 treated groups. BIBN4096 also had no effect on the onset latency of the prolonged TH **(C)** and STH **(D)** sensitization responses or the durations of the TH **(E)** and STH **(F)** sensitization responses following CSD.

## 4. Discussion

The main findings of the study suggest that 1) Acute stimulation of meningeal afferents with high K^+^ concentration promotes a delayed and prolonged increase in their ongoing activity but does not increase their mechanosensitivity, 2) Systemic pretreatment with BIBN4096, a CGRP receptor antagonist, blocks the K^+^ - related prolonged activation of meningeal afferents, and 3) A similar BIBN4096 treatment does not inhibit the prolonged activation or mechanical sensitization of meningeal afferents in response to CSD.

Whether a trigeminovascular reflex involving activity-dependent release of CGRP [16,41] from meningeal afferent peripheral nerve endings contributes to migraine pain remains unclear [57]. Previous studies which examined the effect of acute stimulation of primary afferent neurons that innervate other tissues on the sensitivity of primary afferent nociceptive neurons have yielded conflicting data. For example, studies in monkeys [44] and rats [54] have shown that stimulation of cutaneous nociceptive afferents did not subsequently alter their ongoing activity, mechanosensitivity or heat sensitivity. A study in rabbits, however, reported the development of heat sensitization of nociceptive afferents following antidromic afferent stimulation [23]. Finally, a rat study which employed local capsaicin stimulation to evoke acute excitation of cutaneous afferents documented a delayed and prolonged increase in the afferents’ ongoing activity and mechanosensitivity that were suggested to involve CGRP signaling [39]. Our present findings support a mechanism by which a strong, yet relatively brief (~60 sec), excitation of meningeal afferents at the level of their meningeal RF can give rise to a prolonged increase in their ongoing activity level.

The K^+^ stimulus paradigm used in the current study likely promoted meningeal CGRP release [12,43,58]. Our finding that pretreatment with BIBN4096 abrogated the K^+^ - induced prolonged activation of meningeal afferents suggests that meningeal release of CGRP and its action on non-neuronal meningeal receptors [35] are likely to mediate the prolonged meningeal afferent response. However, given that BIBN4096 was administered systemically (to allow action also on meningeal tissue that remained unexposed in our preparation) we cannot exclude the possibility that inhibition of CGRP receptors localized to the cell body of meningeal afferents in the trigeminal ganglion, which is located outside the blood brain barrier [18,35,14], may have also contributed. Nonetheless, while dural CGRP release is an established phenomenon [12,43,58], evidence supporting CGRP release in the TG, as well as its action receptors localized to the TG cell bodies in vivo is currently lacking.

In addition to the evoked meningeal CGRP release, exogenous stimulation with K^+^ and perhaps CSD-evoked K^+^ release could have also stimulated adrenergic postganglionic autonomic fibers that innervate the meningeal vasculature [56] leading to local release and action of norepinephrine [11]. Such local sympathetic response could have influenced the excitability of meningeal afferents indirectly by promoting the release of prostaglandins [13] or other sensitizing factors from non-neuronal meningeal constituents, such as fibroblasts [68]. However, the earlier finding that norepinephrine signaling inhibits CGRP release from trigeminal afferents [3,27] suggest that activation of meningeal postganglionic sympathetic neurons does not play a role in mediating the prolonged afferent response following K^+^ stimulation.

Our previous finding that local or systemic application of vasodilating doses of CGRP does not activate or sensitize meningeal afferents [37] seems to contradict the current data. The possibility that the levels of CGRP elaborated in response to the K^+^ stimulation in the present study exceeded the concentrations of CGRP employed in our previous work may be entertained. However, previous data suggest that stimulation of meningeal afferents, either electrically or chemically with a mixture of inflammatory mediators or with 50mM K^+^ leads to increased CGRP levels at the picomolar range [12,19,58] - much lower than the micromolar range that failed to influence meningeal afferent responsiveness in our previous study. Differences in the durations of CGRP action between the studies is also an unlikely explanation. In our previous study local CGRP action (measured using changes in local blood flow) lasted for at least 10 minutes when CGRP was applied locally and for ~15 min following systemic application. Thus, the duration of CGRP action in our previous work was likely to be longer than that induced by the release of CGRP in response to the ~1 min excitation of the afferents by the K^+^ stimulus in the present study. As a potential explanation for the seemingly discrepancy between the two studies we propose that the process underlying the K^+^ evoked prolonged afferent activation, which requires CGRP, is multifactorial. The release of CGRP from activated meningeal afferents is likely to be accompanied by the release of other neuropeptides, and potentially by other factors such as glutamate.

These mediators, in turn, could act upon meningeal immune, vascular and Schwann cells, which produce the final algesic mediators that enhance the activity of meningeal afferents. We proposed that CGRP action plays a key part in this cascade of events, through a synergistic effect [5,6], by reinforcing or enhancing the release of other algesic mediators from meningeal non-neuronal cells which express CGRP receptors [35].

In rodent models, CSD is associated with a brief cortical efflux of K^+^ that can reach ~50mM [45,50,66]. Elevated K^+^ in the superficial cortical interstitial fluid could diffuse towards the meninges and is a likely major contributor to the acute activation of meningeal afferents observed in this model [52,63]. Our current data suggest that the prolonged meningeal afferent response that emerges following a brief meningeal K^+^ stimulation occurs with a longer delay and has a shorter duration than that observed following a single CSD episode. These differences, as well as the lack of mechanical sensitization following K^+^ stimulation, suggest that local meningeal K^+^ action, or acute activation of meningeal afferents per se, cannot fully account for the prolonged enhancements in meningeal afferent responses that occur following CSD. The development of mechanical sensitization following CSD [75] but not in response to K^+^ stimulation also points differences between the mechanisms that mediate the prolonged activation and mechanical sensitization of meningeal afferents. This notion is in agreement with our recent data that showed the lack of correlation between CSD-induced afferent activity and mechanosensitization [75] as well as with previous findings from our lab demonstrating the ability of nitric oxide and TNF-α signaling to elicit mechanical sensitization but no activation of meningeal afferents [38,70,72]. It is worth noting nevertheless that CSD is associated with a wave of cortical K^+^ release that spreads throughout the hemicortex, a response that could lead to the acute activation a much larger population of meningeal afferents than following the local K^+^ stimulation used in the current study. Whether a spatial summation of such acute afferent responses plays a role in mediating the prolonged activation and sensitization of meningeal afferents following CSD remains to be determined.

Systemic administration of BIBN4096 is likely to target CGRP receptors primarily outside the central nervous system [14,17]. Our data thus supports the view that acute activation of meningeal CGRP receptor signaling is unlikely to constitute a major contributing factor underlying the prolonged activation and sensitization of meningeal afferents following CSD. Our finding that BIBN4096 partially inhibited the CSD-evoked increase in CBF, a previously recognized CGRP mediated event [8,67] together with its ability to block neurogenic meningeal vasodilatation per se [26,51] also support the claim that acute CGRP-related responses (including its pial vasodilatory effect) are also not major contributing factors that mediated the meningeal nociceptive effect of CSD. The discrepancy between the involvement of CGRP receptor signaling in mediating the K^+^-driven prolonged afferents activation data but not in response to CSD further calls into question the notion that the initial short-lasting cortical efflux of excitatory molecules during the CSD wave, such as K^+^, directly contributes to the prolonged meningeal nociceptive response via acute activation of peripheral CGRP receptors. While the current study suggests the acute CGRP receptor signaling may not mediate the meningeal nociceptive response to CSD, a recent finding suggested that several hours of systemic CGRP sequestering, using anti-CGRP monoclonal antibody (mAb), can inhibit the CSD-induced activation and sensitization of a subpopulation of trigeminal dorsal horn neurons that receive input from meningeal afferents. Such effect however, may be due to an action at the level of the central nerve ending in nucleus caudalis [22,61]. Nonetheless, prolonged inhibition of basal CGRP action, unlike the acute effect of BIBN4096, could have also altered meningeal cellular processing leading to decreased responsiveness of meningeal afferents to CSD in that model.

While CSD can be efficiently induced by a cortical pin-prick stimulus, another common induction method, which has been used in numerous in vivo studies of headache mechanisms, involves epidural application of K^+^ at high concentration (usually 1M). Our finding of acute and prolonged activation of meningeal afferents following epidural K^+^ stimulation at a much lower dose, that does not promote CSD, thus have important implications for the interpretation of previous studies that used K^+^ stimulation to study the effect of CSD on trigeminal/meningeal pain (e.g. [2,59,64,71]). Future studies that employ the CSD paradigm to study related trigeminal responses and their underlying mechanisms thus should be conscientious about the CSD induction method employed. The use of less invasive methods that induce CSD remotely, such as optogenetics [30], which may not influence meningeal afferents responsiveness *per se* may provide a better choice to study mechanisms of CSD-related meningeal nociception and its involvement in migraine headache.

